# GC-MS Profiling and *In vitro* Antibacterial, Anti-Biofilm and Anti-adhesive Activities of *Tamarix ericoides* Rottl. Leaf Extract Against Catheter-Associated Urinary Tract Infectious Agents

**DOI:** 10.1101/2025.01.21.634068

**Authors:** Muhammad Musthafa Poyil, Mohammed H Karrar Alsharif, Mahmoud H. El-Bidawy, Mohammed Saad Alqahtani, Tarig Gasim Mohamed Alarabi, Ahmed Abdullah Albadrani, Alaa Azhari Mohamed Hamid, Abdullah Mohammed Radwan Arafah, Ahmed Abdel Tawab, Saad Alqasem, Ali Al-Gonaim

## Abstract

Catheter-associated urinary tract infection (CAUTI) is one of the most important nosocomial infections among hospitalized patients and causes serious complications due to the development of drug-resistant, biofilms forming strains of microorganisms resulting in treatment challenges. As the chemical catheter-coating agents often fail to prevent biofilm formation, the researchers are looking for compounds of natural origin, and the phytocompounds with multiple modes of action pose as a promising option. Thus, the present study investigated the antibacterial anti-biofilm potentials of the phytocompounds in one of the least explored medicinal plants - *Tamarix ericoides* Rottl. and its leaf methanolic extract was analyzed against *Escherichia coli* – one of the most notorious multidrug-resistant, biofilm-forming uropathogens in CAUTIs. The well-diffusion method showed the antibacterial activity of methanolic leaf extract against *E. coli* and using the microdilution method, the minimal inhibitory concentration (MIC) of the extract against *E. coli* was calculated as 1 mg/ml. GC-MS profiling of methanolic fractions of *T. ericoides* showed the presence of eight important phytochemicals such as diethyl phthalate, ethanol, 2-[2-[(2-ethylhexyl)oxy]ethoxy]-, n-hexadecanoic acid, 9-octadecenoic acid, (E)-, 9,12-octadecadien-1-ol, (Z,Z)-octadecanoic acid, -hydroxy-3-(1,1-dimethylprop-2-enyl) coumarin and Cholestan-3,22,26-triol 16-[2- [formylthio]ethyl]- that are responsible for antibacterial activities. The killing kinetics of *T. ericoides* leaf extract against *E. coli* showed at 1 h. Further, the antibiofilm activity of *T. ericoides* leaf extract against *E. coli* on nonliving surfaces was analyzed and quantified by crystal violet assay. *T. ericoides* leaf extract reduced mature biofilms of *E. coli* by 81%, 85%, and 89 % after treatment with 1X MIC (1 mg/ml), 2X MIC (2 mg/ml), and 3X MIC (3mg/ml) concentrations of the extract respectively. This was further confirmed using SEM analysis wherein biofilm reduction was observed when compared to untreated. The catheter coating with *T. ericoides* leaf extract showed antibacterial activity in the *in vitro* bladder model and was quantified based on colony count. The CLSM reveals the anti-adhesive property of *T. ericoides* leaf extract on the catheter surface which reduced the biofilm formation and biofilm thickness when contacted with *E. coli* cells. Also, 82% of dead cells were observed in the FDA and PI combination. Further, SEM showed the impact of *T. ericoides* leaf extract on *E. coli* cell morphology as the cells displayed damage including cell shrinkage. Furthermore, the leaf extract was found to be non-toxic to normal cells. Based on the findings, the authors recommend further investigation to develop *T. ericoides* leaf extract as a potential catheter coating agent to manage CAUTIs.

## 1. Introduction

Urinary tract infections are among the most commonly encountered nosocomial infectious conditions in hospitalized patients and these infections are mainly attributed to the presence of an indwelling urethral catheter [1], which also cause other complications like ascending UTIs that are vital in urology [2, 3]. The catheters create serious complications like mechanical traumas including symptomatic bacterial infection, urinary leakage, perforation, partial urethral damage, catheter toxicity, hypersensitivity, and anaphylaxis leading to increased lengthy stays and high [4, 5]. The urinary catheters are partially stretchy hollow tubes in structure that are intended to drain the liquid waste from the bladder. Unfortunately, catheters are susceptible to infection because they have direct contact with uropathogens and permit them from the outside environment to the urinary tract which is normally a sterile area damaging the bladder host defense mechanisms [6]. Among the whole device-associated infection, CAUTI represents the second most important infection which ranges up to 40% [7, 8, 9] in hospitalized patients. The significant occurrence of this infection originates from the urinary catheter which allows the opportunistic uropathogens entry through the lumen and makes bacterial adhesion and colonization lead to serious complications such as bladder stones, pyelonephritis, encrustation, bacteriuria, endotoxic shock, and septicemia [10, 11, 12]. Moreover, catheter usage is short or long-term, the patients are easily getting infected and ready to develop biofilms on the inner and outer catheter surfaces giving survival tactics to bacteria [13, 14]. These biofilms are multifaceted distinguished groups encompassing many bacterial associations and produce extracellular polymeric substances that help bacteria to escape from antibiotics and continue to attach on biotic and abiotic surfaces resulting in treatment critical but also improving the development of resistant strains [15, 16]. Several organisms including bacteria and fungi are commonly contaminating the catheters and also, responsible for biofilm development resulting in treatment challenges that lead to high morbidity and mortality. Amongst, one of the most important gram-negative bacteria, *Escherichia coli* is a frequently isolated organism with the ability to form biofilm on catheter surfaces [17, 18]. Generally, CAUTIs are curable but in certain circumstances such as recurrent or inappropriate use of antibiotics increases infection severity [19, 20, 21]. This alarming situation stimulated the search for new antimicrobial drug development with potent antibiofilm as well as anti-adhesive properties to combat CAUTI-causing organisms.

In the past several decades, natural resources have been the better choice for antimicrobial discovery owing to their contribution to pharmaceutics such as antimicrobial, anticancer anti-inflammatory, etc. [22, 23]. The report from WHO says, that most of the global population are seeking plant-based traditional medicine for primary health care [24]. *Tamarix ericoides* is the least studied medicinal plant from the family of Tamaricaceae was commonly used to treat diabetes, gastrointestinal disorders, wounds, and dental problems and also, had anti-inflammatory properties [25, 26]. Based on this indication, our study investigated the antibacterial, anti-biofilm activity of *T. ericoides* methanolic leaf extract against *E. coli* involved in CAUTI.

## 2. Materials and methods

### 2.1. Chemicals and Inoculum Preparation

The chemicals such as FDA, and PI were purchased from Sigma Aldrich, from Sigma Aldrich, Louis, MO, USA and the Mueller-Hinton (MH) broth and rifampicin were purchased from Hi Media, Mumbai, India. The culture *Escherichia coli* was obtained from the American Type Culture Collection (ATCC 25922). The overnight *E. coli* culture grown in MHB adjusted to 10^6^ CFU/ml was used throughout the study and rifampicin and methanol were used as positive and vehicle control respectively.

### 2.2. Preparation of T. ericoides Leaf Methanolic Extract

The methanolic extract of the collected, cleaned air-dried leaves of *T. ericoides* was prepared as per the standard protocol [27]. Briefly, in a new cellulose thimble, about 20 g of fine powder was filled and placed inside the Soxhlet apparatus. Once the methanol was added, the reaction continued for many hours until a clear solution was obtained. The solvent-evaporated product was used for further investigations.

### 2.3. GC-MS Profiling of T. ericoides Leaf Methanolic Extract

To analyze the bioactive fraction from the methanolic extract of *T. ericoides* leaf, GC-MS profiling as per standardized procedure [28]. All the programs like temperature, gas, etc., were initiated, and using a micro syringe, the methanol extracts (1 µl) were injected into the GC-MS. The scanning was continued for 30 mins. Later, the compounds separated were eluted from the column, and with the help of a detector, it was detected using the detector. Each of the peaks in the chromatogram represents an individual molecule present in the extract and it then enters into the mass spectroscopy detector. The identification of compounds was completed by comparing the retention indices and the mass spectra patterns available in the computer library.

### 2.4. Antibacterial Activity of T. ericoides Leaf Methanolic Extract

The methanolic leaf extract of *T. ericoides* antibacterial activity was determined against *E. coli* using the well-diffusion method [29]. Briefly, the well made on a sterile MHA plate swabbed with overnight *E. coli* culture was allowed to receive two different concentrations of methanolic leaf extract and incubated. The zone inhibition around the well indicates the antibacterial activity of *T. ericoides* leaf extract against *E. coli*.

### 2.5. MIC Determination for T. ericoides Leaf Methanolic Extract

To calculate the MIC of methanolic *T. ericoides* leaf extract against *E. coli*, the well diffusion method [30]. Shortly, in MHB broth, 4 mg/ml of *T. ericoides* leaf extract was serially diluted until 0.03mg/ml was received overnight *E. coli* culture. The Optical density of turbidity was measured in all the wells at 600 nm after incubation.

### 2.6. Killing Kinetics of T. ericoides Leaf Methanolic Extract

To determine the killing kinetics of *T. ericoides* leaf extract against *E. coli*, the killing assay was performed [30]. Briefly, *E. coli* (1x10^6^ CFU) overnight culture treated with 1 mg/ml leaf extract was incubated at different time points including 0 h, 1 h, 2 h, 4 h, 6 h, and 12 h. Later, the samples (100 µl) collected from each time point were serially diluted (10-fold), and a spread plate was done for each sample to calculate viable cells by counting CFUs.

### 2.7. Effect of T. ericoides Leaf Extract on Biofilm Formation

To study the influence of *T. ericoides* leaf extract on *E. coli* biofilm formation, the crystal violet assay was used [30]. Briefly, the *E. coli* biofilm formation on a polystyrene plate surface using MHB broth was allowed for 5 days in the presence of various concentrations (4 mg/ ml to 0.03 mg/ml) of *T. ericoides* leaf extract. Later, the formed biofilm washed with phosphate buffer saline (PBS) was fixed with methanol sometimes followed by crystal violet staining. The stained biofilm was detained using acetone and ethanol mixture and the end product was measured at 570 nm.

### 2.8. Effect of T. ericoides Leaf Extract on Mature Biofilm

The qualitative and quantitative *T. ericoides* effect on mature *E. coli* biofilms were studied as per the reported procedure [30]. In brief, the effect of leaf extract on the mature biofilm of *E. coli* was studied qualitatively by allowing the *E. coli* cells to grow on Whatman No.1 filter paper strips for 5 days and treated with 1 mg/ml of leaf extract for 1 h. Later, the attached biofilm on the filter paper strip after washing was fixed with glutaraldehyde dehydrated with varying ethanol gradients and air dried. Then, the gold-coated air-dried filter paper strip was analyzed for SEM images using Supra 55, Carl Zeiss. Similarly, the extract effect on mature *E. coli* biofilm was studied quantitatively using the crystal violet staining method as performed before. In short, on polystyrene surfaces the *E. coli* was allowed to form biofilm for up to 5 days and treated with 1 mg/ml (1X MIC), 2 mg/ml (2X MIC), and 3 mg/ml (3X MIC) for 24 h. The PBS wash was given to remove non-adherent cells followed by crystal staining with methanol-fixed biofilm. The destained final purple color product was read at 570 nm.

### 2.9. Antibacterial Activity of T. ericoides Leaf Extract-Coated Catheters

To investigate the antibacterial effect of *T. ericoides* leaf extract-coated catheter against *E. coli*, an *in vitro* bladder model [32] analysis was performed. The air-dried sterile small catheter tube coated with leaf extract was placed over the prepared MHA plates swabbed with overnight *E. coli* culture and incubated for clear zone inhibition around the tube which demonstrates the antibacterial activity of *T. ericoides* leaf extract coated catheter against *E. coli*.

### 2.10. Quantification of Bacterial Load on the Bladder Model

To quantify the bacterial load from *T. ericoides* leaf extract-coated catheter against *E. coli,* the viable bacterial count method was followed as mentioned before [33]. In short, the extract-coated and non-coating small sterile catheter tube was allowed into contact with *E. coli* culture for 24 h. Later, the catheter tube was transferred to a fresh centrifuge tube and the tube was shaken vigorously to dislodge the attached cells followed by 10-fold serial dilution for both samples. Then, 100 µl of treated and untreated samples were used to calculate viable cell count based on CFU count, and another 100 µl samples from both samples were used to measure turbidity at 600 nm to calculate growth percentage.

### 2.11. Visualization of Bladder Model

To visualize the non-adhesive property of *T. ericoides* leaf extract-coated catheters against *E. coli,* confocal microscopy was used [33]. Briefly, a small sterile catheter tube coated with leaf extract was permitted to form biofilm on the catheter surface by submerging the catheter tube into *E. coli* culture containing broth for five days followed by PBS wash. The washed tube was stained with fluorescein diacetate (FDA, 40 µl from 5 mg/ml) for10 mins followed by propidium iodide (PI, 20 µl from 1 mg/ml) for 5 mins and the images were observed using CLSM to calculate live/dead cell percentage.

### 2.12. Effect of T. ericoides Leaf Methanolic Extract on Cell Morphology

To understand the effect of *T. ericoides* leaf extract on *E. coli* cell morphology, scanning electron microscopy analysis was employed [31]. Briefly, *E. coli* cells grown on Whatmann No.1 filter paper were treated with 1mg/ ml of leaf extract for 1h followed by PBS wash to remove unattached cells. The glutaraldehyde fixation was done for washed paper strips and dehydration with ethanol gradient. The strips were observed after gold coating for morphological changes and the changes were captured using SEM (Supra 55, Carl Zeiss, Wetzlar, Germany).

### 2.13. Cytotoxicity of T. ericoides Leaf Methanolic Extract

To study the *T. ericoides* leaf extract’s cytotoxic effect on L_929_ cells, the MTT assay [30] was conducted. In short, the cells grown in Dulbecco’s Modified Eagles Medium (DMEM) with 10% fetal bovine serum were reacted with various concentrations of *T. ericoides* leaf extract (0.5, 1, 2, 3, and 4 mg/ml) for 24 h followed by formazan formation was observed after the addition of MTT solution. Later, the formazan crystals dissolved were read at 570 nm after adding DMSO, and the cell viability percentage was calculated using the following formula.

Cell viability percentage= [(Treated cells OD)/ (Untreated cells OD)] ×100

### 2.14. Statistical Analysis

The mean and standard deviations were calculated for all the experiments like MIC determination, extract effect on biofilm formation, and biofilm eradication. Statistical significance was carried out for quantification of bacterial load and live dead assay using student *t*-test and also, the significance was predicted when p-value ≤0.05.

## 3. Results

### 3.1. GC-MS Profiling of T. ericoides Leaf Methanolic Extract

The GC-MS profiling of extracted leaf extract of *T. ericoides* analyzed for the identification of various compounds present in the methanolic fraction is presented in Figure 1. The compound structure was analyzed based on the fragmentation pattern of mass and the spectral data comparison with chemical profiles located in the National Institute of Standards and Technology (NIST) library. As seen in Figure, the chromatogram of eight important peaks represents eight individual compounds such as Diethyl Phthalate, Ethanol, 2-[2-[(2-ethylhexyl)oxy]ethoxy]-, n-Hexadecanoic acid, 9-Octadecenoic acid, (E)-, 9,12-Octadecadien-1-ol, (Z,Z)-Octadecanoic acid, - Hydroxy-3-(1,1-dimethylprop-2-enyl) coumarin and Cholestan-3,22,26-triol 16-[2- [formylthio]ethyl]- were observed in methanolic leaf extract of *T. ericoides*. The retention time, peak area, percentage peak area, height percentage, identified compound name, and structure are presented in Table 1.

**Figure 1.**
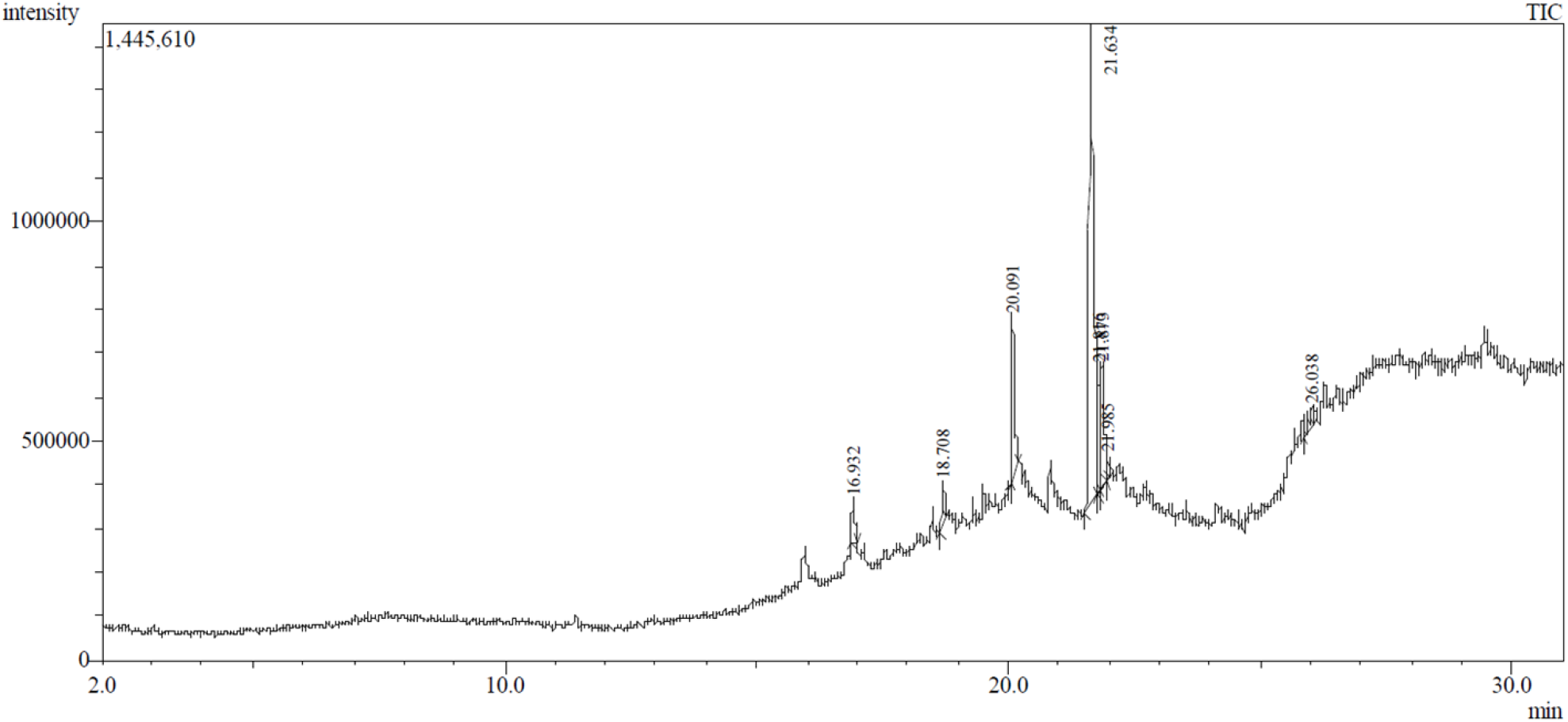
GC-MS chromatogram of *T. ericoides* leaf methanolic extract.

**Table 1.**
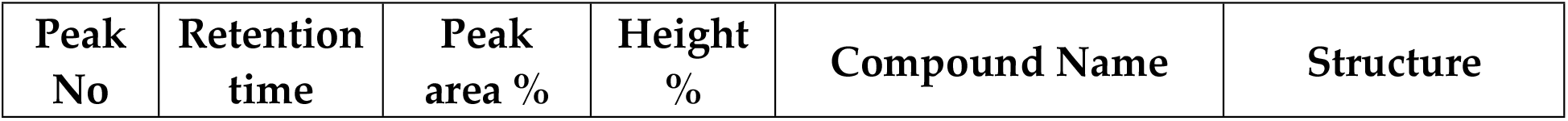
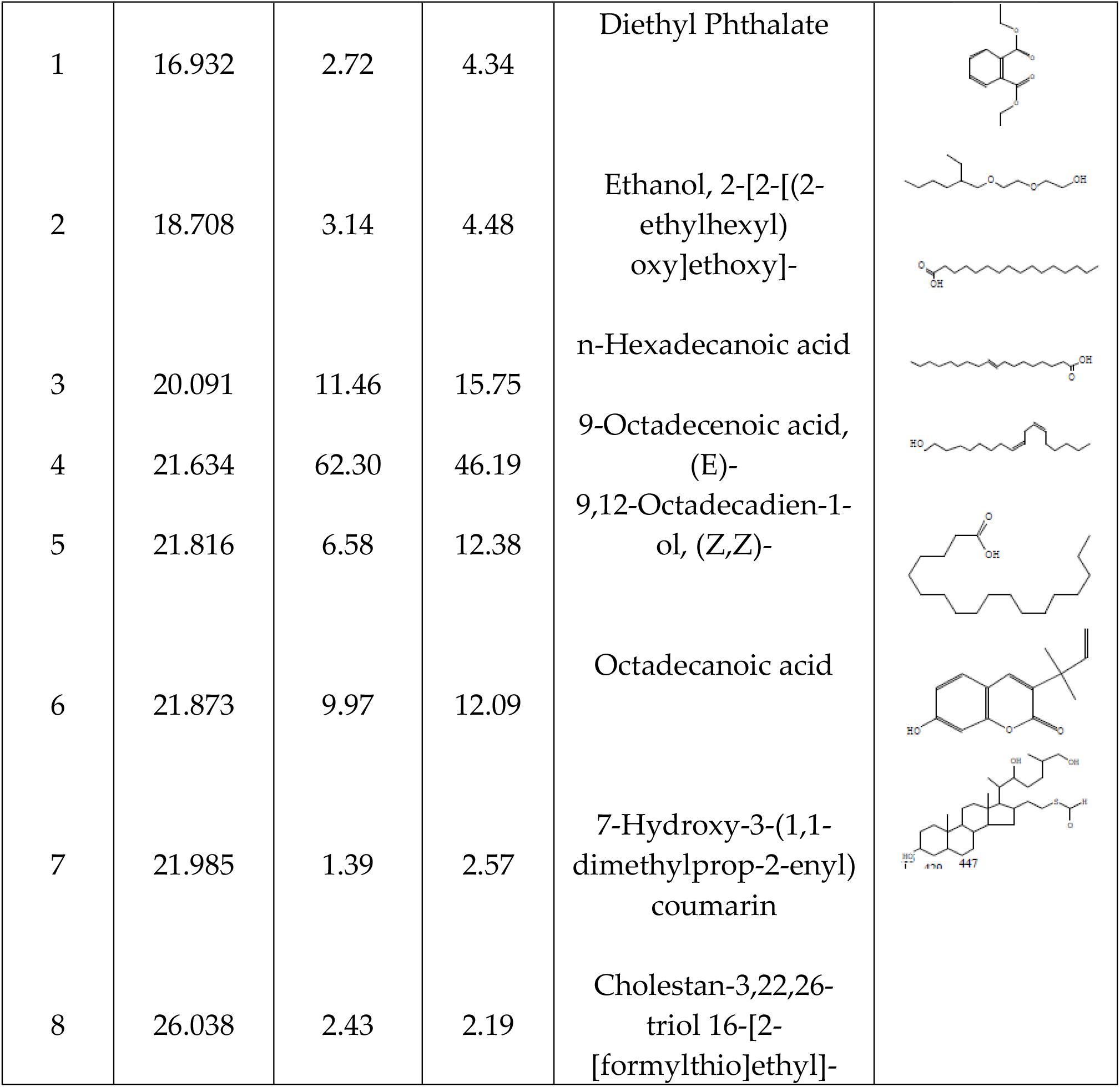
Phytochemicals obtained from methanolic fractions of *T. ericoides* leaf using GC-MS.

| Peak No | Retention time | Peak area % | Height % | Compound Name | Structure |
| --- | --- | --- | --- | --- | --- |
| 1 | 16.932 | 2.72 | 4.34 | Diethyl Phthalate |  |
| 2 | 18.708 | 3.14 | 4.48 | Ethanol, 2-[2-[(2-ethylhexyl)oxy]ethoxy]- |  |
| 3 | 20.091 | 11.46 | 15.75 | n-Hexadecanoic acid |  |
| 4 | 21.634 | 62.30 | 46.19 | 9-Octadecenoic acid, (E)- |  |
| 5 | 21.816 | 6.58 | 12.38 | 9,12-Octadecadien-1-ol, (Z,Z)- |  |
| 6 | 21.873 | 9.97 | 12.09 | Octadecanoic acid |  |
| 7 | 21.985 | 1.39 | 2.57 | 7-Hydroxy-3-(1,1-dimethylprop-2-enyl) coumarin |  |
| 8 | 26.038 | 2.43 | 2.19 | Cholestan-3,22,26-triol 16-[2-[formylthio]ethyl]- |  |

### 3.2. Antibacterial Activity of T. ericoides Leaf Methanolic Extract

The antibacterial activity of methanolic *T. ericoides* leaf extract was confirmed against *E. coli* and the observed growth inhibition around the well using the well-diffusion method is presented in Figure 2. As seen in the figure, the increasing zone size was evidenced in dual concentrations (2 mg and 3 mg/ml) of leaf extract demonstrating the antibacterial activity was concentrations dependant.

**Figure 2.**
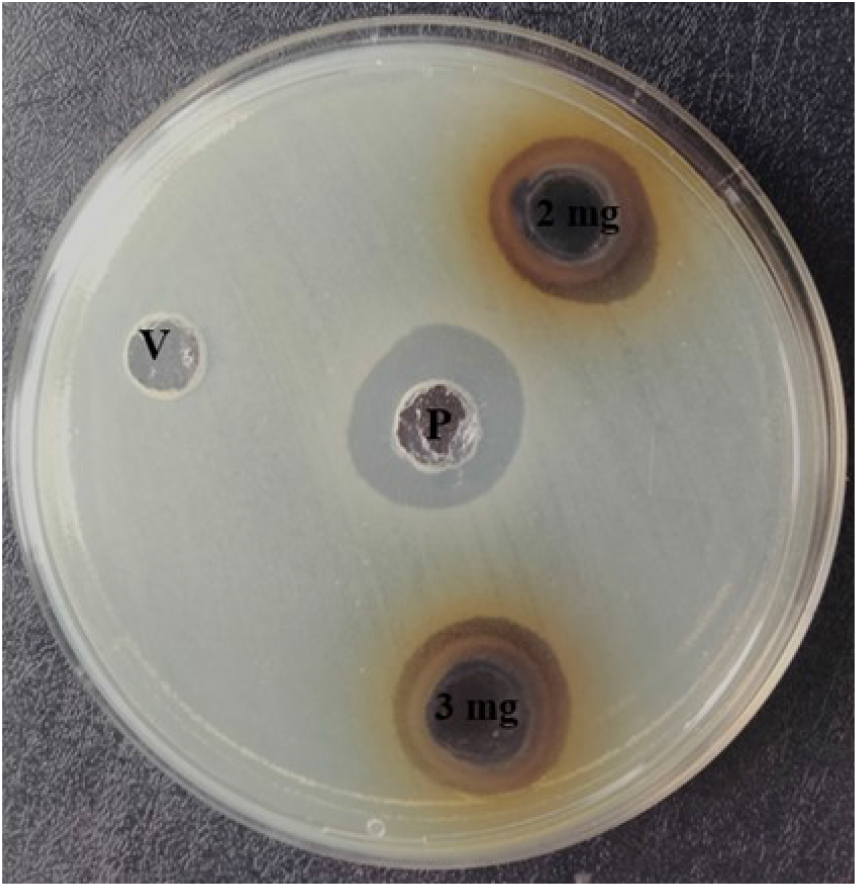
*T. ericoides* leaf extract antibacterial activity against *E. coli.* Note: V-vehicle control and P- positive control

### 3.3. MIC Determination for T. ericoides Leaf Methanolic Extract

The lowest concentration required to inhibit *E. coli* growth calculated using the micro-dilution method is presented in Figure 3. The MIC of *T. ericoides* leaf methanolic extract against *E. coli* growth was 1 mg/ml.

**Figure 3.**
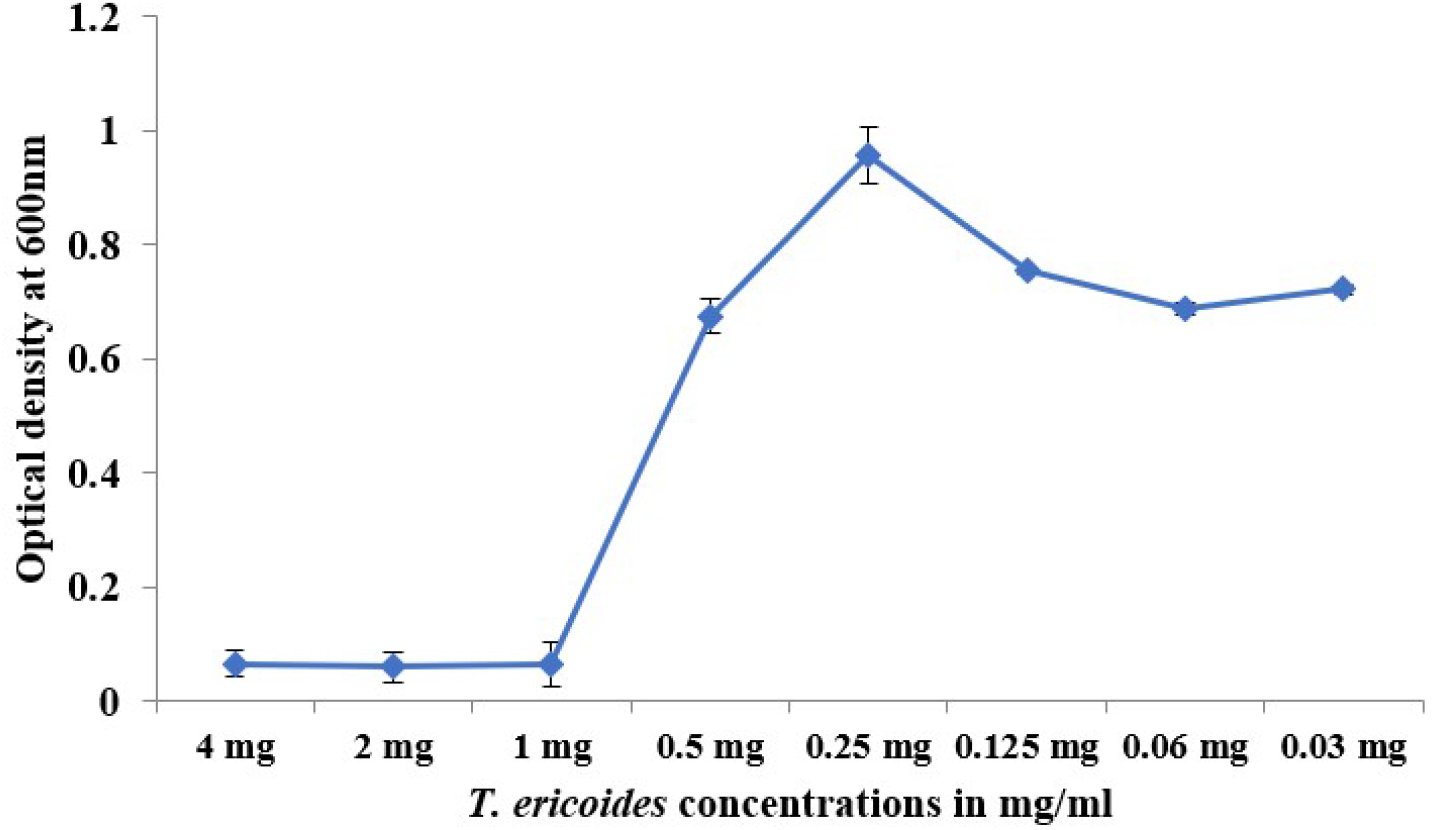
MIC determination for *T. ericoides* leaf methanolic extract

### 3.4. Killing Kinetics of T. Ericoides Leaf Extract

The investigated killing kinetics and calculated growth inhibitory effect of *T. ericoides* leaf extract against *E. coli* using a killing assay are represented in Figure 4. The figure shows that the leaf extract-treated *E. coli* cells exhibited no life after 1 h but cells without treatment showed more viable cells.

**Figure 4.**
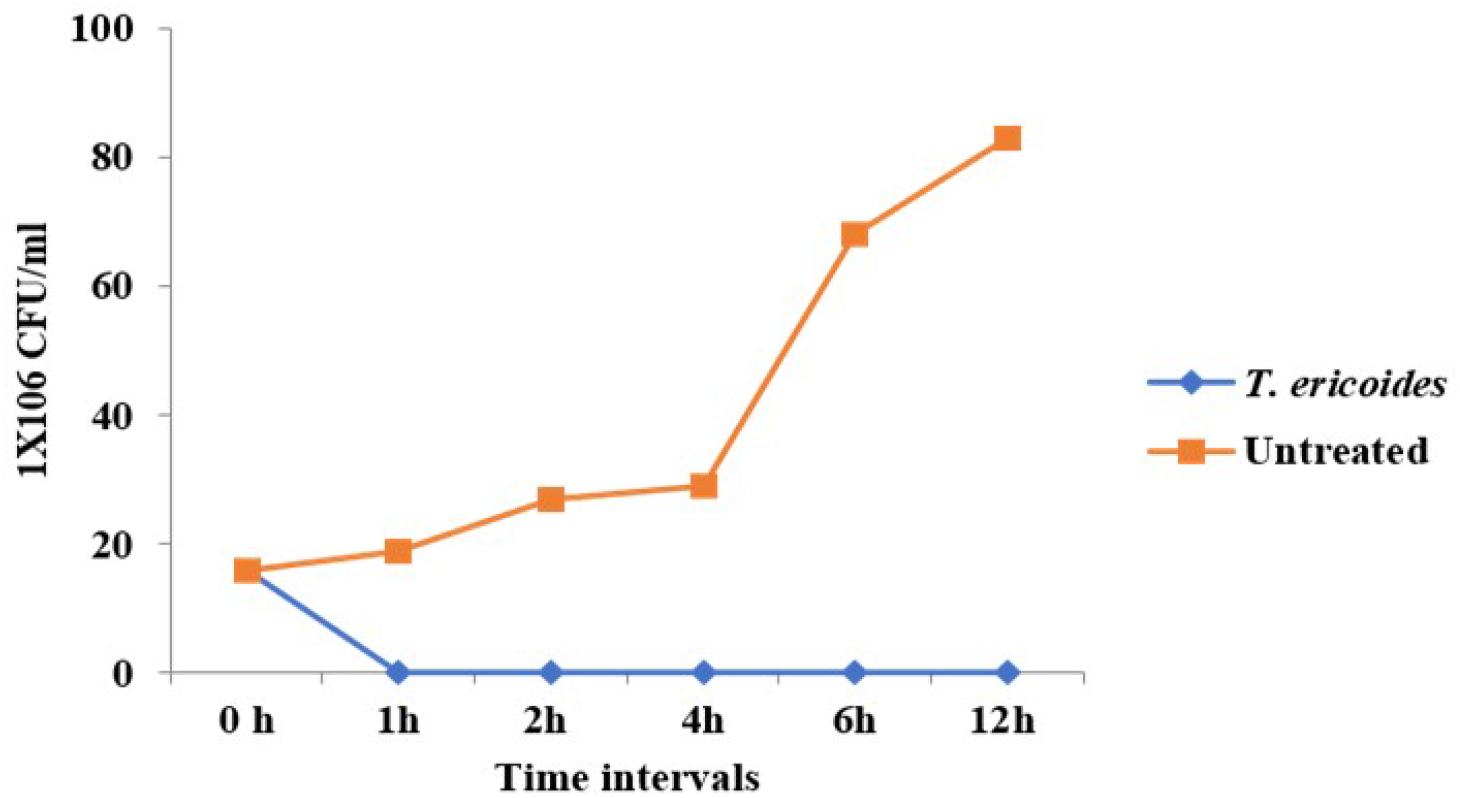
Killing kinetics of *T. ericoides* leaf extract showed no live *E. coli* cells after 1 h treatment.

### 3.5. Effect of T. ericoides Leaf Methanolic Extract on Biofilm Formation

The effect of *T. ericoides* leaf extract on *E. coli* biofilm formation was studied quantitatively using crystal violet assay and the biofilm formation percentage after leaf extract treatment is displayed in Figure 5. The Figure verified the biofilm inhibiting capacity of varying ranges of leaf extract concentrations against *E. coli.* Moreover, the biofilm formation was inhibited until the MIC of leaf extract and the gradual increase of biofilm formation was noted after its MIC represents the traces of extract can also able to slow down the biofilm formation.

**Figure 5.**
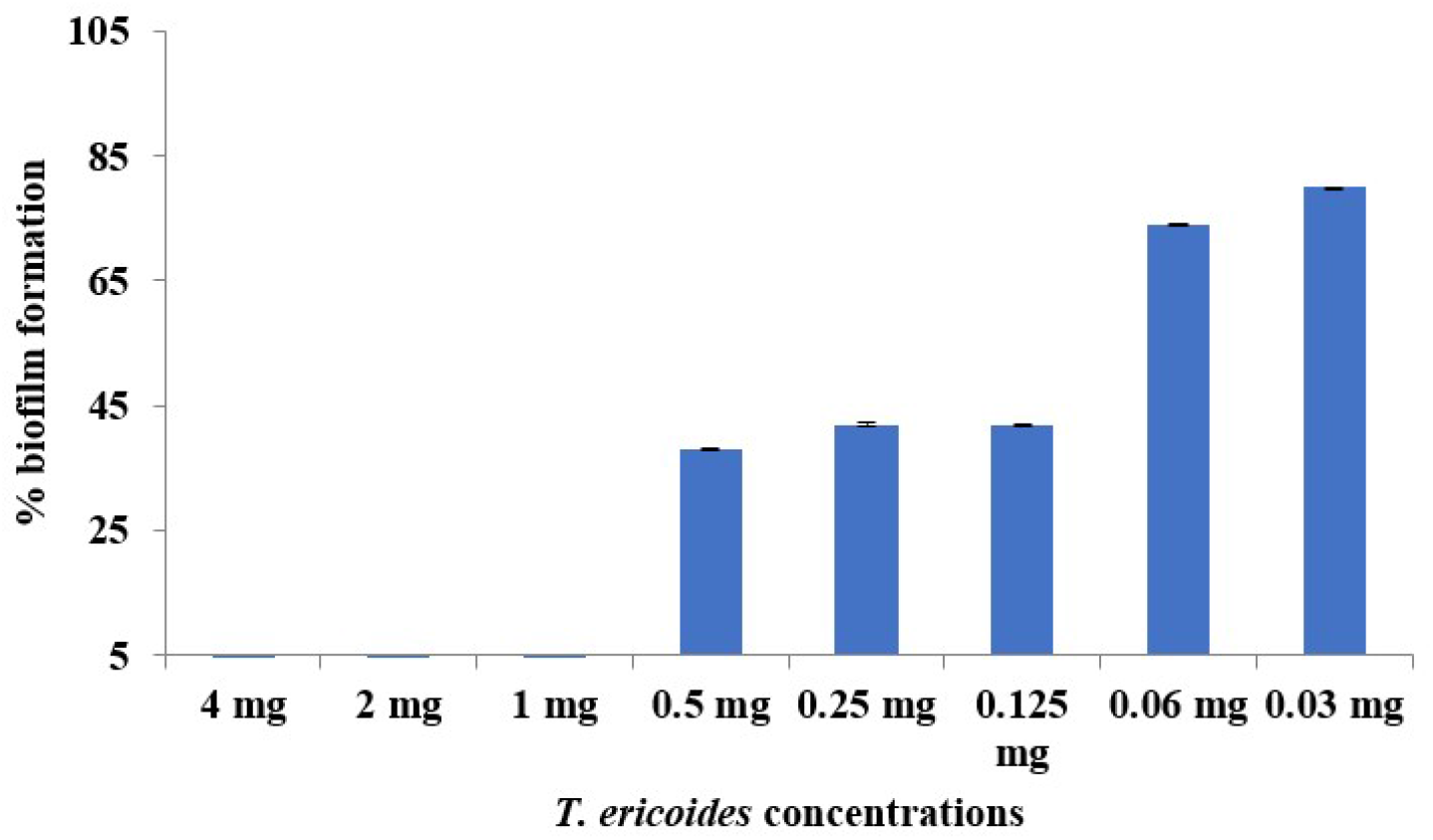
*T. ericoides* leaf extract influence on *E. coli* biofilm formation was investigated along with various leaf extract concentrations that showed biofilm inhibition until 1 mg/ml.

### 3.6. Effect of T. ericoides Leaf Extract on Mature Biofilm

*T. ericoides* leaf extract effect on *E. coli* mature biofilm was studied qualitatively on cellulose matrices and the pictorial representation of biofilm eradication after leaf extract treatment is presented in Figure 6. The Figure shows an SEM image of the minimal number of adherent cells on the cellulose matrix after treatment with leaf extract resulting in biofilm eradication when compared to the untreated matrix wherein abundant adherent cells were noticed on the matrix thus no biofilm eradication. In addition, three different *T. ericoides* leaf extract concentrations treated with *E. coli* mature biofilms quantified using the crystal violet method are presented in Figure 7. The Figure shows the percentage of biofilm eradication after *T. ericoides* 1X, 2X, and 3X MIC concentrations treatment which reduced 81%, 85%, and 89 % of *E. coli* mature biofilms respectively.

**Figure 6.**
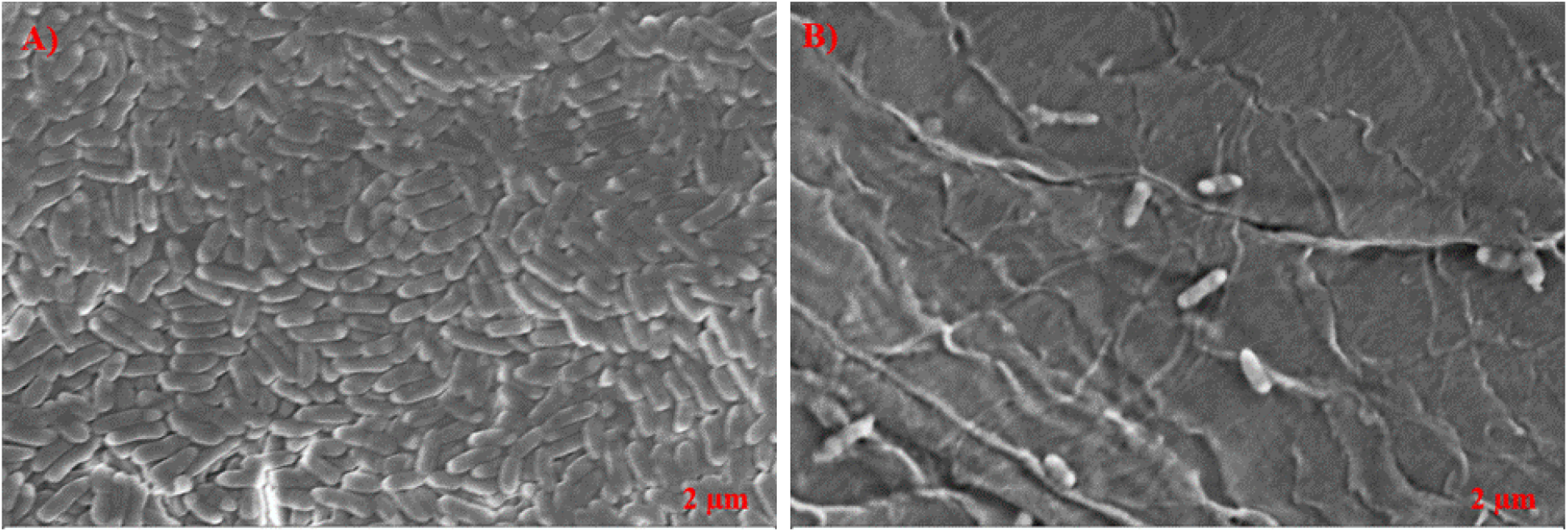
A qualitative study of *T. ericoides* leaf extract impact on biofilm eradication after treatment. A) SEM image of viable cells adherent on cellulose matrix B) Leaf extract treatment reduced the attached cells on the matrix was evidenced by SEM. Scale bar- 2µm.

**Figure 7.**
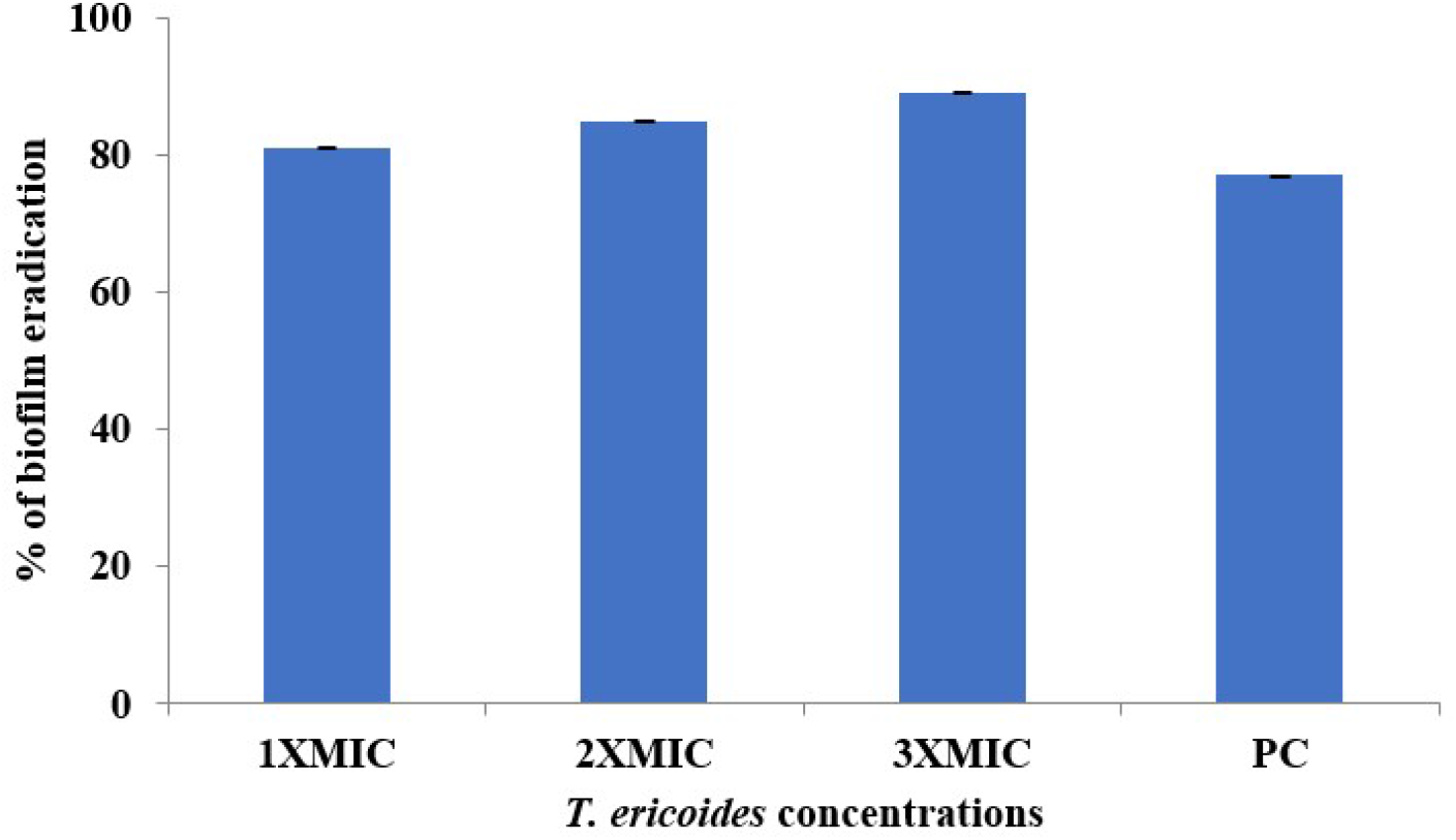
Quantitative representation of biofilm eradication after treatment with three concentrations of *T. ericoides* leaf extract.

### 3.7. Antibacterial Activity of T. ericoides Leaf Extract-Coated Catheters

The catheter tube coated with *T. ericoides* leaf extract was investigated for antibacterial activity against *E. coli* using an *in vitro* bladder model and the zone formation around the catheter tube is displayed in Figure 8. The zone formation indicated the antibacterial activity of the catheter tube coating with leaf extract. In addition, the bacterial load was quantified from a catheter tube coated with *T. ericoides* leaf extract using the colony counting method, and the calculated growth percentage of leaf extract is displayed in Figure 9 A-C which shows the least number of viable cells were counted in coated catheter tube when compared to uncoated catheter tube and also, the calculated growth percentage was 14% in coated catheter tube against *E. coli* represents the anti-adhesive property of *T. ericoides* leaf extract.

**Figure 8.**
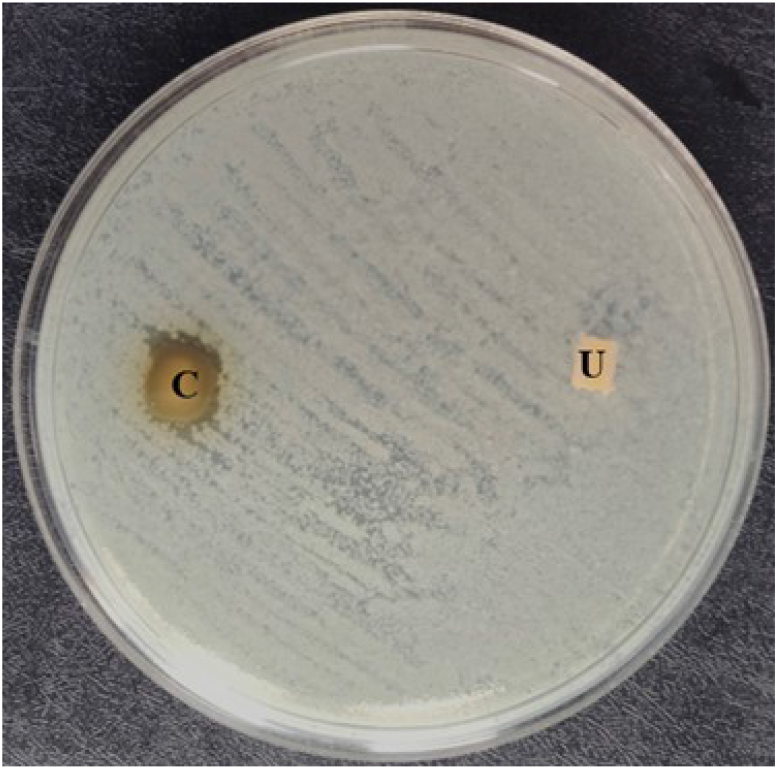
The catheter coated with *T. ericoides* leaf extract antibacterial activity was investigated against *E. coli* and showed zone development around the catheter tube.

**Figure 9.**
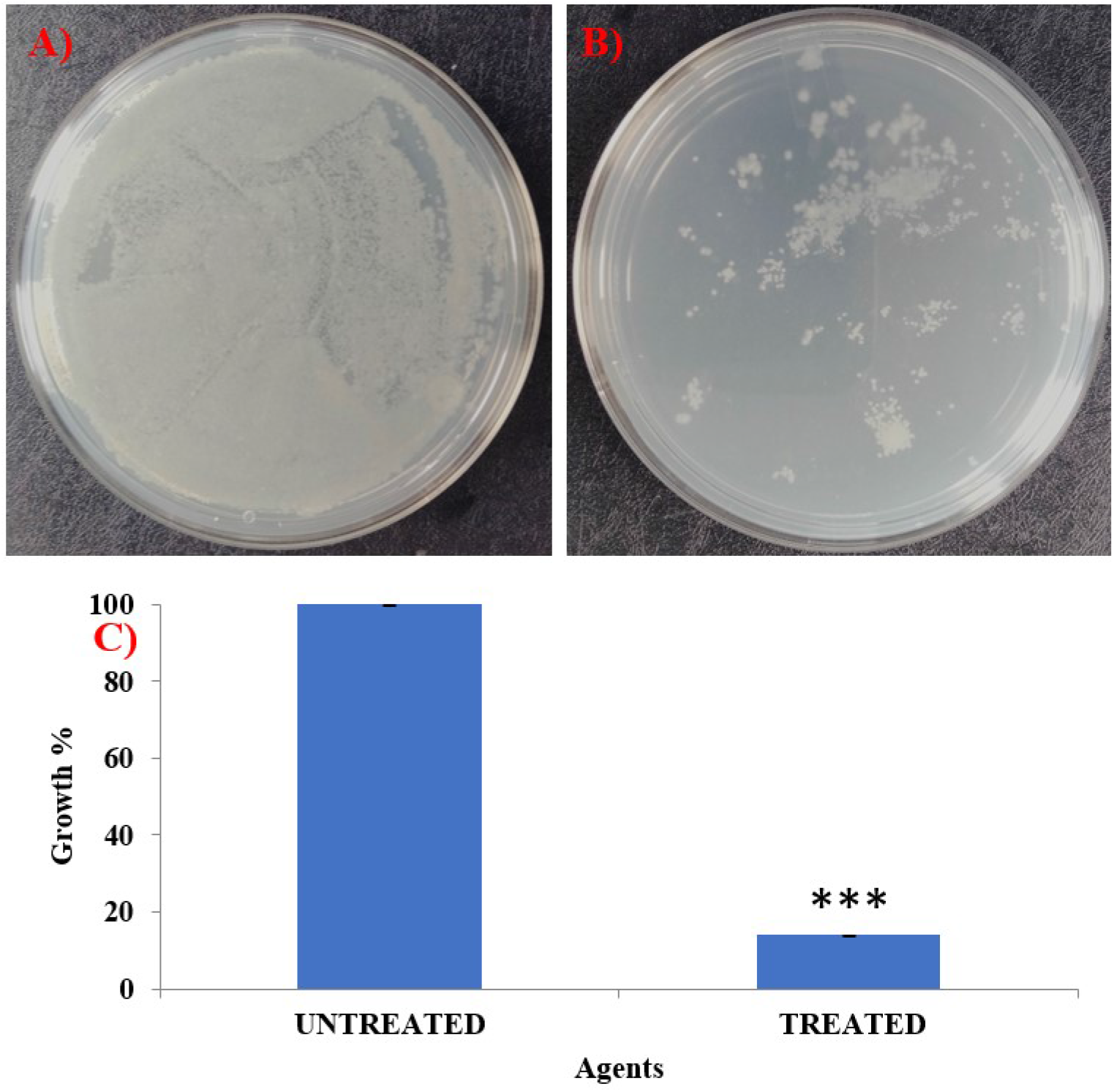
Bacterial load quantified from coated and uncoated catheter tube against *E. coli*. A) More number of CFUs found in the uncoated catheter tube B) Leaf extract coated catheter tube revealed the less CFUs C) Graph denotes the growth percentage after coating the leaf extract. *** Highly significant

### 3.8. Visualization of Bladder Model

The anti-adhesive property of *T. ericoides* leaf extract-coated catheter tube was visualized after five days of contact with *E. coli* cells through CLSM and the calculated percentage of live/dead cells is mentioned in Figure 10A-D. The catheter tube without coating was contacted with *E. coli* cells for 5 days after being stained with fluorescein diacetate (FDA, binds to live cells which emit green fluorescence) and propidium iodide (PI, binds to DNA of membrane damaged cells emit red fluorescence) mentioned in Figure 10A. Figure 10B denotes the three-dimensional view of E. coli biofilm thickness (25µm) on a catheter tube. As seen in Figure 10C, high red fluorescence was observed on the coated catheter which represents the leaf extract damaged the cell membrane and binds to DNA. As indicated in Figure 10D, catheters coated with leaf extract have the antiadhesive property against *E. coli* cells on the catheter surface which was evidenced through biofilm thickness reduction in three-dimensional structure (14 µm) when compared to uncoated catheters. Moreover, the FDA and PI combination provides a pictorial representation of live/ dead on the catheter surface, and also, the percentage of live/ dead cells calculated based on FDA and PI in combination is represented in Figure 11. *T. ericoides* leaf extract-coated catheter displayed 82% of dead cells after treatment.

**Figure 10.**
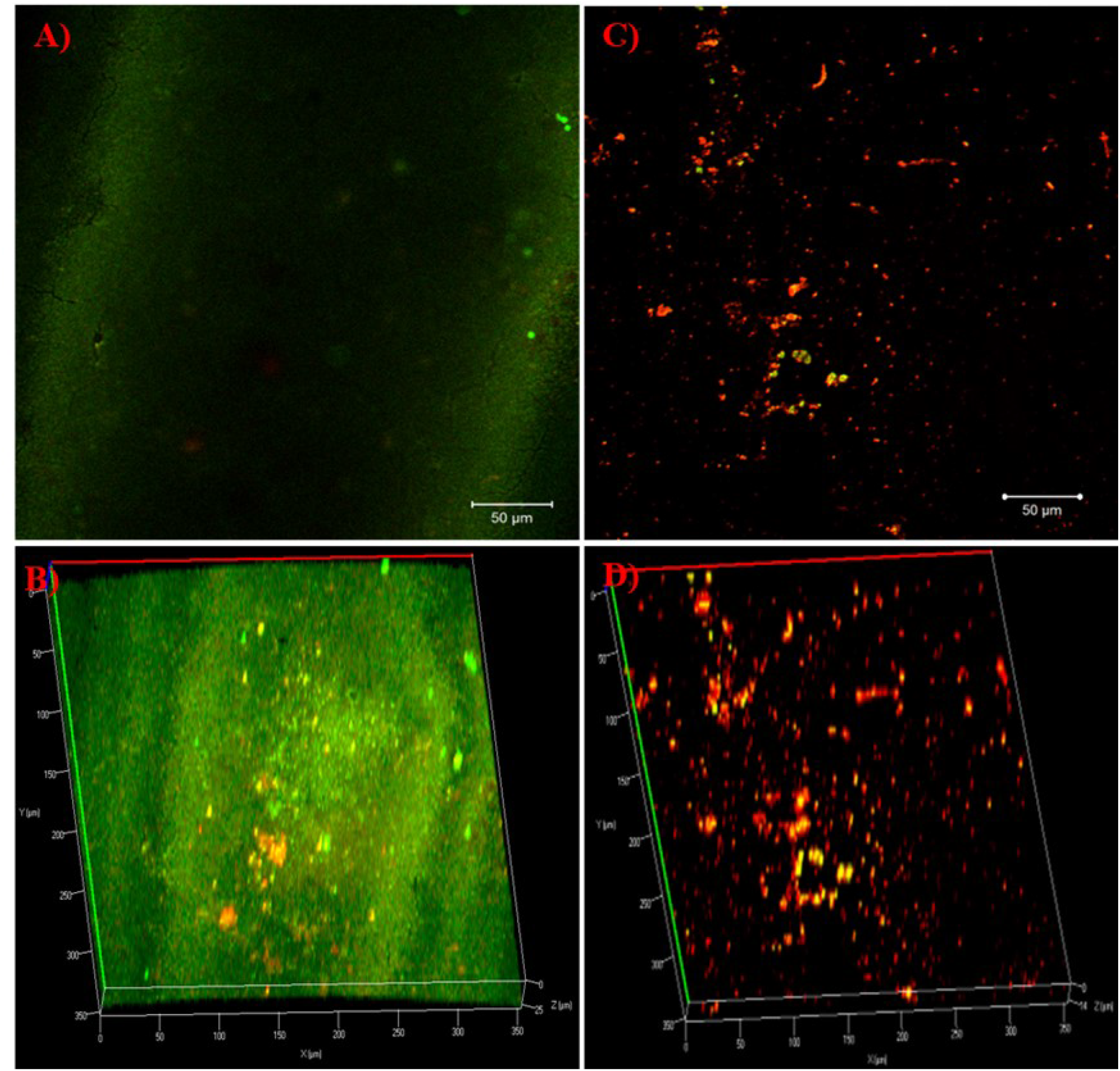
Confocal Microscopy of the catheter tube. A) Uncoated catheter tube reveals biofilm formation on their surface after 5 days of contact with *E. coli* B) Three-dimensional view of biofilm formation on catheter surface which represents 25 µm thickness C) Catheter coated with *T. ericoides* showed biofilm eradication D) Three-dimensional structure revealed the reduction in biofilm thickness till 14 µm.

**Figure 11.**
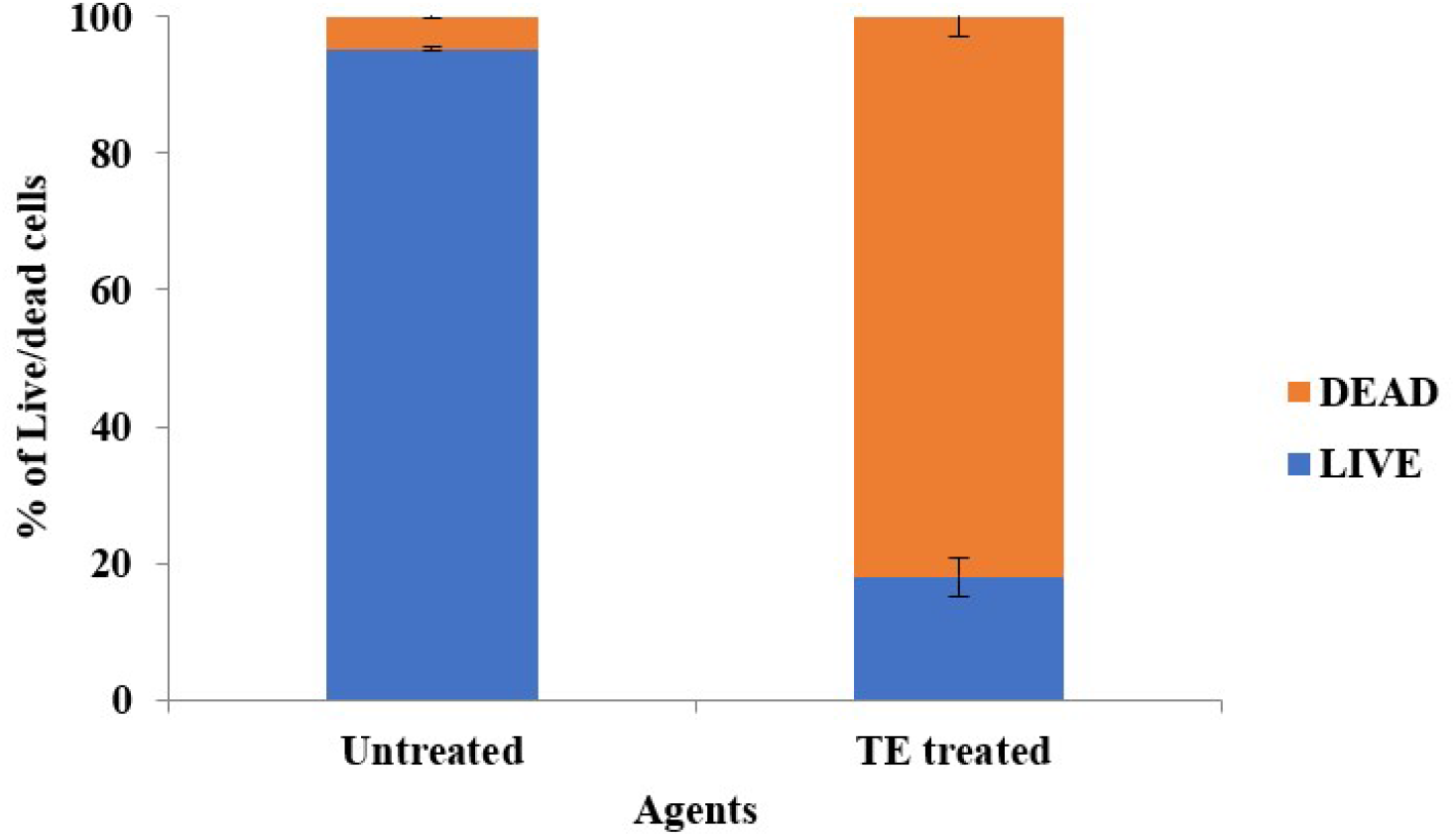
Live and dead *E. coli* cells percentage from catheter tube coated with leaf extract and uncoated tube was calculated and showed 82% of dead cells after treatment.

### 3.9. Effect of T. ericoides Leaf Extract on Cell Morphology

The SEM showed the changes that occurred in *E. coli* cell morphology during contact with the *T. ericoides* leaf extract for 1 h examined and the attained images are presented in Figure 12. As monitored in Figure, the cell shrinkage was noted due to internal cell leakage when treated with leaf extract; whereas undamaged cell structure was observed on untreated *E. coli* cells.

**Figure 12.**
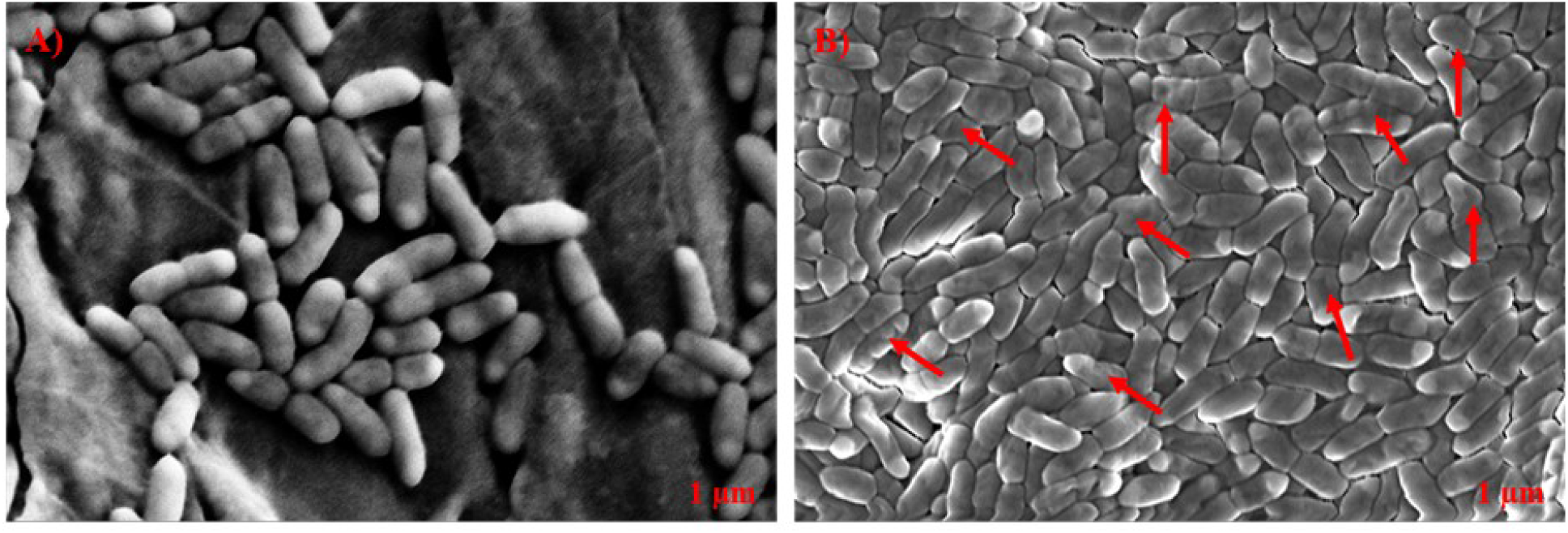
*T. ericoides* leaf extract effect on *E. coli* cell morphology. A) Untreated E. coli cells with undamaged morphology B) Red arrow indicates cell shrinkage after *T. ericoides* leaf extract.

### 3.10. Cytotoxicity of T. ericoides Leaf Extract

*T. ericoides* leaf extract cytotoxicity studied on L_929_ cells after treatment with varying concentrations is presented in Figure 13. The cell viability percentage after treatment was calculated for all the concentrations and the cell viability observed at 1 mg/ml was 91% proving that the leaf extract was not toxic to L_929_ cells compared to untreated cells.

**Figure 13.**
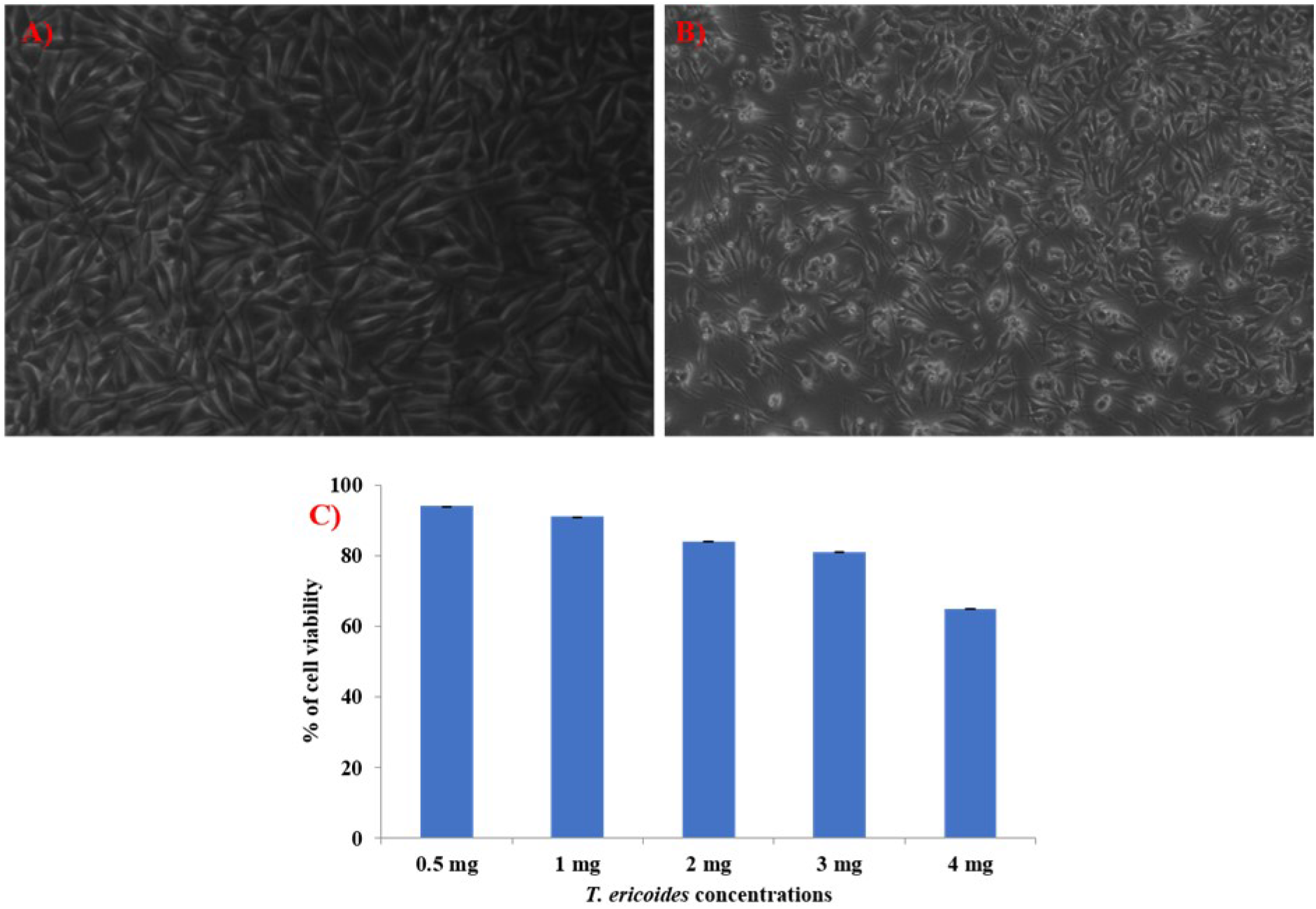
*T. ericoides* leaf extract cytotoxicity was studied on L_929_ cells A) Untreated cells B) Cells treated with Extract C) The cell viability percentage calculated after varying concentrations denotes leaf extract was not cytotoxic to normal cells.

## 4. Discussion

Device-associated infections, particularly CAUTIs are a significant medical complication that occurs in hospitalized patients for various reasons. Generally, CAUTI is curable but sometimes the treatment process is ineffective due to biofilm-forming organisms such as *E. coli* an important uropathogen, and also, overuse of antibiotics makes treatment challenges as well as resistant strains development. This dreadful situation prompted us to find an alternative antibacterial agent immediately to fight against CAUTI-causing organisms. Hence, our study investigated the antibacterial activity of methanolic *T. ericoides* leaf extract against *E. coli* and found antibacterial activity using very low concentrations. Our study was supported by a recent work wherein *Tamarix ericoides* Rottler, an unfamiliar medicinal plant was extracted by various solvents like methanol, ethanol, aqueous solutions petroleum ether, etc., and studied the antimicrobial activity against *Bacillus subtilis, Salmonella typhi, E. coli,* and *C. albicans* and found the potent antimicrobial activity in methanol extract when compared to other solvents [34]. Similarly, the antimicrobial activity of *T. nilotica* (Ehrenb) Bunge from the Tamaricaceae family was investigated against *Klebsiella pneumoniae,* and strong antibacterial activity was observed in n-butanol fractions [35]. Likewise, numerous reports investigated *T. aphylla* from Saudi Arabia’s medicinal plant’s antimicrobial activity against various human pathogens. The antimicrobial activity of both methanolic and ethanolic leaf extract against *E. coli*, *B. subtilis*, *S. typhi*, *S. aureus*, *Aspergillus flavus*, and *C. albicans* showed potent activity and suggested that the solvents had no differences in their MICs against test pathogens [36,37,38]. In addition, different species such as *T. gallica and T. ramosissima* from the Tamaricaceae family were investigated against human pathogens and found that they had strong activity against gram-negative organisms rather than gram-positive and fungi owing to the biomolecules present in the extracts [39,40,41,42]. Similarly, the antimicrobial activity was further investigated by killing kinetics against *E. coli* to determine the dose of leaf extract for various studies.

The phytochemicals present in the methanolic fraction of *T. ericoides* leaf extract were investigated to find out the phytochemicals such as diethyl Phthalate, Ethanol, 2-[2-[(2-ethylhexyl)oxy]ethoxy]-, n-Hexadecanoic acid, 9- Octadecenoic acid, (E)-, 9,12-Octadecadien-1-ol, (Z,Z)- Octadecanoic acid, -Hydroxy- 3-(1,1-dimethylprop-2-enyl) coumarin and Cholestan-3,22,26-triol 16-[2- [formylthio]ethyl]- in methanolic leaf extract of *T. ericoides* and these are responsible for antibacterial activity against *E. coli*. Many reports say diethyl Phthalate, 9- Octadecenoic acid, and coumarin showed antimicrobial activity against many important including *Staphylococcus aureus, Pseudomonas aeruginosa*, *Candida albicans,* [43,44,45,46,47,48,49]. The phytochemicals profiling of 6,10,14-trimethyl-2- pentadecanone, dodecanoic acid, and octadecane are major compounds along with many minor compounds were reported in hexane fractions of *T. aphylla* [50]. The three major compounds such as hispidulin, isorhamnetin, and cirsimaritin were identified in *Tamarix ramosissima* bark extract and they were found to inhibit the formation of 2-amino-1-methyl-6-phenylimidazo[4,5-*b*] pyridine suggesting their great potential beneficial effects on human health [51]. All the results represent different species that have different phytochemicals for different applications.

In addition, *T. ericoides* leaf extract antibiofilm activity was investigated against *E. coli* an important uropathogen that can able to form biofilm formation on the catheter surface and makes treatment ineffective. Normally, the uropathogens enter into the catheter from the exterior environment resulting development of biofilm including many stages such as attachment, colony formation, and maturation which makes treatment less effective [52,53]. Hence, our study focused on each stage of biofilm formation to prevent biofilm formation, the *T. ericoides* inhibited the biofilm formation of *E. coli* on non-living surfaces the antibiofilm activity was proved. The antibiofilm activity was quantified by investigating the activity on mature biofilms and eradicating the biofilms after treatment. In support of this, electron microscopy revealed the *T. ericoides* effect on *E. coli* biofilms by eradicating or reducing biofilm after treatment. Similarly, a recent study evaluated dichloromethane (DCM) and ethyl acetate (EtOAc) fractions of *T. nilotica* antifungal activity against clinical isolates of *Candida albicans* and demonstrated antifungal activity with the least inhibitory concentration of 64-256 and 128-1024 µg/mL, respectively. The SEM examination revealed reduced or decreased biofilm formation DCM fraction treatment suggesting *T. nilotica* can be an important source antifungal agent against *C. albicans* [54].

Further, the formed biofilms can able to block the catheter and protect the bacteria from host defense and antibiotic treatment resulting in antibiotics failure to eliminate the bacteria present in the biofilms [55,56]. To prevent biofilm formation on the catheter’s inner and outer surfaces, the coating catheter with any antimicrobial agent is an excellent method [57]. Henceforth, our study proved the *T. ericoides* extract-coated catheter antimicrobial activity against test pathogens. It was further supported by the quantification of bacteria from a catheter coated with extract revealed minimal viable cells of *E. coli* when compared to an uncoated catheter which suggests that the catheter coated with *T. ericoides* extract prevents biofilm formation. To confirm the biofilm inhibition on the catheter surface, the coated and uncoated catheter contact with *E. coli* was visualized by staining with FDA and PI revealing the development of structured biofilms on the uncoated catheter surface and reduced biofilms on the coated catheter surface which strongly suggest that, the catheter coated with *T. ericoides* extract is efficient in preventing biofilm formation on the catheter surface. Our findings were supported by many groups wherein abundant antimicrobial agents including antibiotic combinations, fosfomycin, silver nanoparticles, polymer, and zinc oxide coated with catheter exposed the robust antimicrobial activity against *S. aureus, E. faecalis*, *K. pneumoniae,* and *E. coli* [58,59,60,61,62].

Apart from that, when *E. coli* contact with *T. ericoides* extract creates cell damage representing weakened cells and also osmotically unstable cells which creates cells more permeable to internal components resulting in leakage of inner components and finally cell death which was evidenced through SEM. Finally, our goal is intended for human use, hence, the *T. ericoides* extract toxicity was checked for safety purposes which showed no toxic effect on normal cells suggesting that, the methanolic *T. ericoides* extract can be a useful antibacterial agent for CAUTI infection.

## 5. Conclusion

The methanolic leaf extract of *T. ericoides* was investigated against CAUTI-causing uropathogens*, E. coli,* and exhibited antibacterial activity at low concentrations due to the phytochemicals present in the extract which was evidenced in GC-MS analysis. The *T. ericoides* showed killing kinetics against *E. coli* and the antibiofilm activity was proved by inhibiting biofilm formation and also, eradicating mature biofilms after treatment on non-living surfaces and it was further confirmed by SEM analysis. The antiadhesive property of *T. ericoides* leaf extract was studied using an *in vitro* bladder model wherein the antibacterial activity was determined and it was further quantified based on colony count. The CLSM reveals the visual effect of *T. ericoides* leaf extract on the catheter surface which suggests the antiadhesive property of leaf extract against *E. coli. T. ericoides* leaf extract was not toxic to normal cells. Based on the above findings, the *T. ericoides* leaf extract can be a potent coating antibacterial agent against *E. coli*.

## Acknowledgments

The authors extend their appreciation to Prince Sattam bin Abdulaziz University for funding this research work through the project number (PSAU/ 2024/03/29416)

